# Assessing male reproductive investment in Papaveraceae using flow cytometry reveals lineage-specific trajectories to pollen-to-ovule ratio reduction

**DOI:** 10.1101/2024.08.23.609364

**Authors:** Jurriaan M. de Vos, Yannick Woudstra, Ilia J. Leitch, Oriane Hidalgo

## Abstract

Male reproductive investment, in particular pollen production, is a crucial and ecologically relevant component of a plant’s phenotype and reproductive success. Its evolutionary trajectory, however, remains understudied, partly due to a lack of convenient methods to assess it. We developed a protocol for pollen quantification by flow cytometry and applied it to 107 flowers from 38 Papaveraceae species differing widely in floral traits (e.g., floral symmetry, stamen number), pollination syndromes (e.g., wind and insect pollination) and reproductive systems (e.g., degree of autogamy). We phylogenetically tested whether pollen number evolved in association with ovule, carpel, stamen and flower numbers per inflorescence, and if there were interacting effects between floral symmetry and/or self-compatibility with pollen and ovule production.

Compared to manual counts, results using flow cytometry were similar, but much faster to obtain and more precise. Pollen and ovule numbers per flower varied > 39,000x and > 550x, respectively, among species. Pollen production correlated positively with ovule, carpel and stamen numbers. Lineage-specific trajectories to pollen-to-ovule ratio reduction (to values < 300) are observed. One involved increased female investment in ruderal species belonging to the subfamily Papaveroideae, while the other occurs through decreased male investment and is associated with the evolution of floral traits towards greater specialisation. The impact of reproductive systems on male and female investment is limited to ovule production in non-actinomorphic flowers.

Taken together, these results revealed that the evolutionary associations between reproductive systems, floral traits, and pollen and ovule production are lineage-specific. Given the profound contrasts at the subfamily level of Papaveraceae, broader surveys across the diversity of flowering plants are clearly needed to better understand factors driving the evolution of reproductive investment. Such studies will certainly be facilitated by our new high-throughput pollen counting method outlined here.

## Introduction

The vast diversity of angiosperm flowers is commonly interpreted as a response to the immobility of plants and the need for vectors to transport pollen from the androecium of one plant to the female organs of another (Barrett, 2010). Plants have evolved diverse reproductive systems (e.g., self-compatibility) and strategies (e.g., wind pollination, such as in many grasses and trees, versus animal pollination, found in poppies and other “visually attractive” flowers) that are reflected in a variety of male and female investment (pollen and ovule number, respectively, and pollen-to-ovule ratio – P/O; Cruden, 1977, 2000; Harder & Johnson, 2023) and concomitant floral diversity. In addition to its importance for ensuring successful plant reproduction, investment in the production of pollen is crucial for sustaining the diversity and abundance of pollen harvesting floral visitors, since it represents a major food reward (along with nectar), for pollination services in plant-insect mutualistic relationships (Müller *et al*., 2006). However, our knowledge of how male reproductive investment varies throughout angiosperm lineages remains very fragmentary and incomplete (Erbar & Langlotz, 2005; Barrett & Harder, 2017).

At least in part, the relatively poor understanding of male investment is due to limited convenient methods for quantifying pollen production. For example, manual pollen counts, using haemocytometers, are often prohibitively time-consuming and imprecise (Bechar *et al*., 1997; Dafni, Kevan & Husband, 2005); electronic particle counters (e.g., Coulter Counter; Beckman Coulter, Fullerton, California, USA; Elzone Micromeritics Instrument Corporation, Norcross, Georgia, USA) are often unavailable; while methods relying on digital imaging require calibration or extensive sample preparation to function reliably (Costa & Yang, 2009; Staedler *et al*., 2018). Such issues make broad surveys challenging. Thus, there is an urgent need for simple, accurate, precise, cost-effective, high-throughput methods that enable large-scale pollen quantification. Flow cytometers have become increasingly available in many laboratories due to the wide range of versatile applications in the plant sciences (Kron, Suda & Husband, 2007), including the analysis of cytogenetic and reproductive traits (e.g., Pellicer, Powell & Leitch, 2021; Kron *et al*., 2021). This study aims to provide a simple and cost-effective protocol for counting pollen using flow cytometry in order to investigate drivers of male reproductive investment.

To demonstrate the effectiveness of this approach, we carried out the first family-level study of pollen and ovule production in the Papaveraceae s.l. (APG, 2009; 44 genera, c. 820 species; Ranunculales). Papaveraceae is particularly suited as a study system, since it displays an exceptionally wide spectrum of floral diversity with four major, angiosperm-wide evolutionary trends that are expected to impact pollen number and P/O. First, they present marked variation in stamen number (from four to several hundred; Ronse De Craene & Smets, 1995) indicating shifts to polyandry (i.e., flowers with more than twice as many stamens as petals; Sokoloff *et al*., 2007) and suggesting a broad range of pollen production. Secondly, they display a striking diversity of floral symmetry, with a progressive transition from actinomorphy to zygomorphy, including an unusual intermediate disymmetric state (Hidalgo & Gleissberg, 2010; Hidalgo, Bartholmes & Gleissberg, 2012; Zhao *et al*., 2018). Floral symmetry, which evolves in concert with floral complexity (Krishna & Keasar, 2018) is a crucial component in plant-pollinator interaction, thought to influence flower attractiveness, pollinator fidelity and pollen transfer efficiency (Citerne *et al*., 2010; Culbert & Forrest, 2016). Factors affecting P/O, such as pollen production and outcrossing rate, are impacted by floral symmetry (Cruden, 1977). However, there is still little understanding of the specific links between pollen production, ovule production and the suite of traits involved in floral symmetry changes. Thirdly, Papaveraceae offer one of the best-studied self-incompatibility systems from a molecular perspective (*Papaver rhoea*s L.; Wang *et al*., 2019, and references therein). Importantly, there are numerous transitions between self-compatibility (SC) and self-incompatibility (SI; Bilinski & Kohn, 2012; Table S1), one of the most common trends across flowering plants and of major importance for floral- and lineage diversification (Igic, Lande & Kohn, 2008; Barrett, 2010; de Vos *et al*., 2014a; Barrett & Harder, 2017). Pollen-to-ovule ratios have long been known to be affected by the amount of autogamy (Cruden, 1977), but the specific relationships between pollen production, ovule production, and plant mating types remain poorly understood (Barrett & Harder, 2017). Fourthly, flowers of Papaveraceae range from solitary (e.g., *Romneya coulteri* Harv.) to inflorescences with up to hundreds of flowers [e.g., *Macleaya cordata* (Willd.) R.Br.], they are either determinate or indeterminate in growth, and the flowering sequence is either acropetal or basipetal (Hidalgo & Gleissberg, 2010). The diversity of inflorescence architectures in the family appears tightly linked to transitions in floral symmetry (Hidalgo & Gleissberg, 2010) as well as mode of pollination (Lidén, 1986; Blattner & Kadereit, 1999). Increasing evidence stresses the importance of integrating inflorescence data for a thorough understanding of reproductive investment (Harder *et al*., 2004; Harder & Prusinkiewicz, 2013; Liao & Harder, 2014). Finally, these four major trends may be related (e.g., Jabbour, Damerval & Nadot, 2008), suggesting their effects on pollen and ovule numbers should be studied in concert. Thus, Papaveraceae offers an unusually wide array of possibilities for asking fundamental questions on the evolution, ecology and biology of plant reproduction, that require rapid and accurate quantification of pollen.

Here, we disentangle the impacts of floral traits –specifically those involved in polyandry, floral symmetry, and self-incompatibility– on pollen number, ovule number and their P/O ratio, enabled by developing an accurate, high-throughput method to quantify pollen number using flow cytometry. We test for correlated evolution of floral and inflorescence traits in a phylogenetic framework and determine whether pollen and ovule production are intrinsically more variable than P/O. Our results indicate that effects of floral traits, including self-compatibility, on pollen number, ovule number, and P/O may be lineage-specific. Such findings highlight the need for additional broad surveys of reproductive traits in other families to further understand the extent to which they impact reproductive success –something that the development of the new method described here could help to achieve.

## Materials and methods

### Plant material

Fresh plant material was either collected in the field or obtained from plants growing in cultivation at botanic gardens (Table S1). Where sufficient material was available, five flower buds were analysed, collected from one or several individuals. Compatibility type was scored from primary literature (Table S1). For each flower sampled for pollen analysis we recorded its position on the inflorescence and the number of flowers per inflorescence together with the number of stamens, carpels, and ovules.

### Pollen collection

Pollen samples were obtained by removing mature, non-dehisced anthers from floral buds and storing them in open 1.5 ml microcentrifuge tubes in a closed box with silica gel to dry for at least one week before analysing. For flowers with more than six stamens, five stamens were taken from the different pseudo-whorls and the obtained pollen count would then be adjusted based on the total number of stamens in the flower. For flowers with six or less stamens, the whole androecium was sampled.

### Pollen counts by flow cytometry

Lycopod spore tablets were used as a standard for pollen counting (9666 +/- 671 spores per capsule; Batch 3862; obtained from the Department of Quaternary Geology, Lund University, Lund, Sweden; https://www.geology.lu.se/services/pollen-tablets). One Lycopod tablet was placed in each flow cytometry tube, suspended in 1 mL of 1N HCl and vortexed until it had completely dissolved, suspending its individual spores. The dry anthers were crushed with a dissection needle in their microcentrifuge tube, suspended in 500 μL of ‘General purpose buffer’ (GPB; Loureiro *et al*., 2007) supplemented with 3% PVP-40, vortexed, sonicated for 1 min at 30 kHz (SFE 590/1 ultrasonicator, Ultrawave Limited, Cardiff, UK), vortexed again and briefly spun in a microcentrifuge to collect any liquid from the lid of the tube. The pollen suspension was then filtered through a 150 μm nylon mesh (Partec, Münster, Germany) into the Lycopod spore suspension. Anthers were rinsed with an additional 500 μL of GPB, vortexed and the suspension was filtered into the Lycopod spore suspension. Propidium iodide was then added to the sample (to a final concentration of 50 μg·mL^−1^) to stain the pollen exine and tubes were placed on ice for 30 min. Each sample was analysed using a CyFlow SL3 flow cytometer (Partec, Münster, Germany) fitted with a 100-mW green solid-state laser (Cobolt Samba, Solna, Sweden). Data were acquired using the logarithmic mode (FloMax software v2.7, Partec). Samples were run until a minimum of 200 pollen grains and 200 spores was reached, or until 1000 spores were obtained when pollen count was very low. The total number of pollen grains per sample (representing the pollen production of the number of stamens counted) was calculated by multiplying the ratio between counted pollen and spore particles with the number of lycopod spores in the tablet (i.e., 9666). This value represented the number of pollen grains per flower for species with six or less stamens, and had to be adjusted to the total number of stamens for species with more than six stamens. Measurements were performed in triplicate for each sample, taking the mean as our estimate of pollen production.

### Manual pollen counts

To compare the flow cytometry method with the common approach of manual counting (Dafni, Kevan & Husband, 2005), we estimated the number of pollen grains for one or two flowers per species with a haemocytometer (Todd & Vansell, 1942) using the standard protocol described in Carleial *et al*. (2017).

### Phylogeny reconstruction

We gathered GenBank nucleotide sequences (Table S2) for two chloroplast markers (*matK* and *rbcL)* for all sampled species (or a closely related substitute) and three outgroup species: *Thalictrum javanicum* Blume (Ranunculaceae), *Euptelea polyandra* Siebold & Zucc. (Eupteleaceae) and *Magnolia grandiflora* L. (Magnoliaceae). We produced sequence alignments using MAFFT in Geneious 8.1.9 (Biomatters Ltd., Auckland, New Zealand) with manual corrections preceding concatenation, and reconstructed the maximum-likelihood phylogeny using the RAxML-HPC BlackBox tool in the CIPRES portal (Miller, Pfeiffer & Schwartz, 2010) with default settings and 100 bootstrap replicates.

### Precision and accuracy of flow cytometry

To assess the accuracy of our pollen counting protocol, we tested whether the estimated number of pollen grains obtained using flow cytometry (Table S3) differed systematically from that obtained with a haemocytometer (Table S4) using a phylogenetic paired two-tailed t-test using the R phytools package (Revell, 2012). This test accounted for standard errors (SE) of all counts. Each paired sample consisted of the mean and associated standard error for typically 3-4 flowers measured with flow cytometry, and the mean among typically five estimates from a single flower (collected on the same day from the same individual or accession) and its standard error measured with a haemocytometer. We thus account for the within-individual variation of pollen number per flower, and imprecision of both pollen counting methods. This test included 29 species. Similarly, to test whether the precision of flow cytometry exceeded that of manual counts, we compared standard errors of counts obtained with both methods using a phylogenetic paired t-test.

### Evolutionary diversification of pollen and ovule numbers

We illustrate one application of our method, by revealing the correlated evolution of pollen number per flower with stamen, carpel, and ovule number per flower, P/O, and flower number. For this, we computed bivariate phylomorphospace plots (Revell, 2012), that display phylogenetically linked, ln-transformed species-mean values with ancestral states estimated using Brownian Motion. The ancestral states were based on the maximum-likelihood phylogram transformed with the maximum likelihood λ-value (Pagel, 1994). Because λ-values were close to 1 (see results), Brownian Motion was an appropriate model of evolution for ancestral character state reconstruction. For each trait-pair, we tested for evolutionary association using Pearson correlation coefficients of phylogenetic independent contrasts (Felsenstein, 1985). These analyses included 38 species (Table S1). We then tested the effects of floral symmetry, self-compatibility, and their interaction, on pollen production per inflorescence (pollen per flower multiplied by the number of flowers in an inflorescence) using phylogenetic generalised least squares (PGLS) with co-estimated λ-parameters, as recommended (Revell, 2010), implemented in the R-package caper (Orme *et al*., 2018). Similarly, we tested their effects on ovule production per plant and on P/O. For PGLS, it was necessary to score zygomorphic and disymmetric species as “non-actinomorphic”, due to limited sample size; these models therefore included 31 species. We compared the relative variability of pollen number, ovule number and P/O as the standard deviation of natural log-transformed species means, testing whether intrinsic variability differed across traits using an F-test (Lewontin, 1966).

## Results

### Pollen count using flow cytometry: performance and reliability

The raw output from the flow cytometer showed that Papaveraceae pollen could be distinguished from Lycopod spores (e.g., Fig. 1A-C), which enabled us to gate the pollen and spore area and determine the particle number in each. While Lycopod spore area appeared consistently circular on the Relative fluorescence versus side scatter (SSC) flow histogram plots, pollen area was more variable in shape (e.g., compare shape of red and blue areas in Figs. 1A-C), indicating heterogeneity in PI staining and/or fluorescence between the pollen particles within the same flower. The flow cytometry results were consistent with the manual counts, but marginally more precise (Figs 1D-E). Species-mean pollen counts did not differ between methods (t_26_ = -0.273, p = 0.787; Fig 1D), but the number of particles processed in each flow cytometry run (mean 630 spores and 424 pollen grains; Table S3) greatly exceeded that of manual counts. Also, the technical error of flow cytometry, expressed as standard deviation among three runs of the same sample, was much smaller (typically less than 10% of the pollen count; Table S3) than with the manual count (Table S4). Likewise, the standard error of the species mean was marginally significantly lower than that of manual counting (phylogenetic paired t-test, t_26_ = -1.792, p = 0.0847; Fig. 1E).

**Figure 1.**
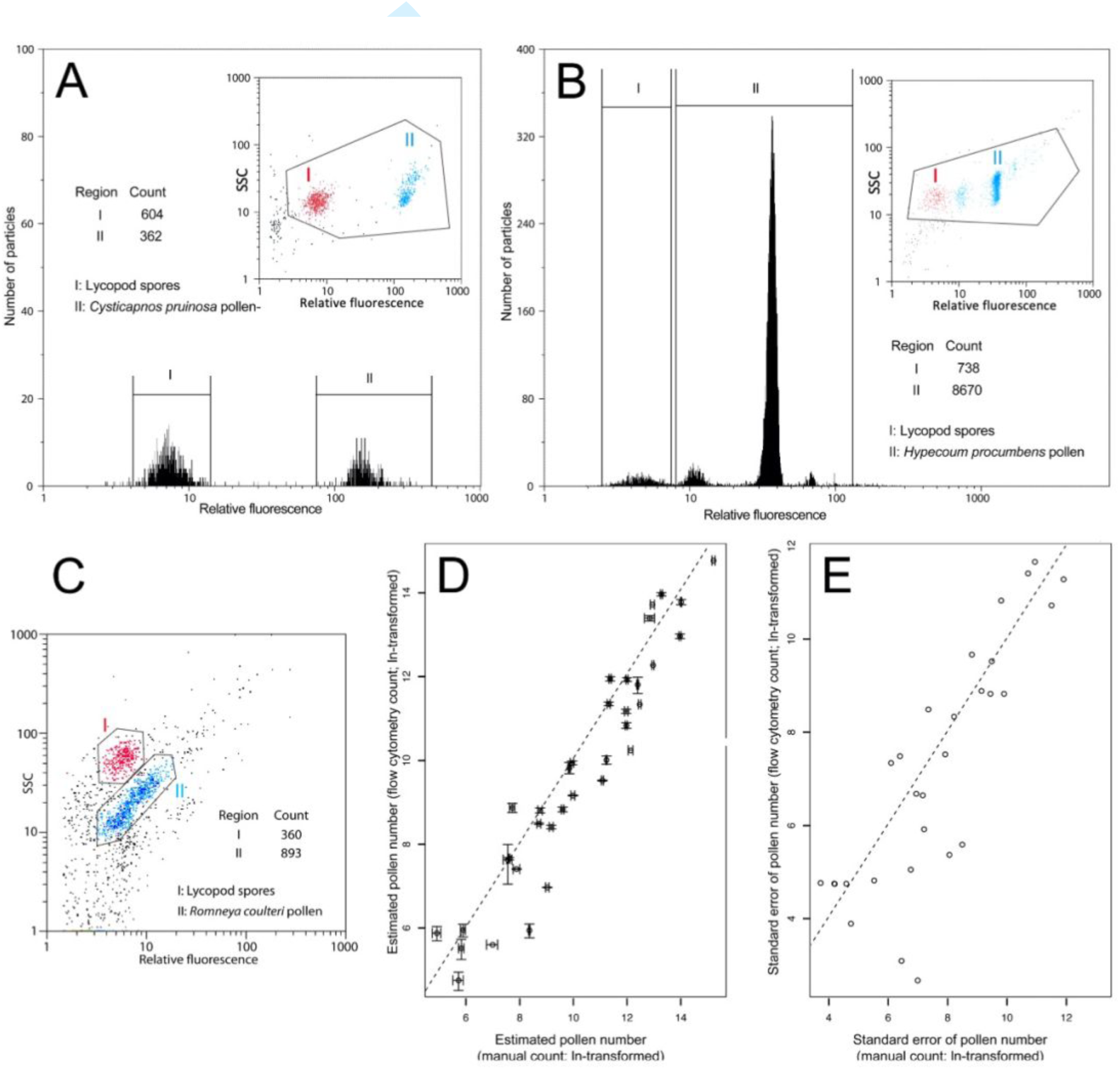
Measurement of pollen grain number using flow cytometry and its comparison to manual counts. (A-C) Panels depict raw output histograms from the flow cytometer for *Cysticapnos pruinosa* (Bernh.) Lidén (A), *Hypecoum procumbens* L. (B), and *Romneya coulteri* Harv. (C), respectively. Gated areas are indicated. (D), Scatterplot showing the relationship between the estimated pollen numbers per sample using manual counts and flow cytometry, with error bars extending 1 SE. (E) Comparison between the standard errors of manual and flow cytometric counts. The dotted lines in panels D and E indicate the relation y = x.

### Variation of pollen number and P/O in relation to the evolutionary diversification of Papaveraceae flowers

Mean pollen number per flower varied > 39,000-fold among Papaveraceae species, ranging from 63.5 (SE 13.25) in *Fumaria capreolata* L. to 2,494,654 (SE 333,327) in *Romneya coulteri* (Table S3). Mean ovule number per flower varied three orders of magnitude among species, from one in *Fumaria* L. and *Platycapnos* Bernh. to 3,420 in *Papaver somniferum* L. P/O varied similarly from 20 in *Papaver dubium* L. to 37,033 in *Macleaya cordata* (Table S1). The relative variability of pollen and ovule number per flower greatly exceeded that of P/O, indicating that a wide range of pollen and ovule numbers contributed to similar P/O values (Fig. 2). Specifically, the variance of log-transformed species-means were 7.48 for pollen number, 5.58 for ovule number and 2.18 for P/O, these being significantly different for pollen vs. P/O (F_df=37_ = 3.43, p < 0.001) and for ovules vs. P/O (F_df=37_ = 2.56, p < 0.005). The most striking examples illustrating this result include *Romneya coulteri* and *Pteridophyllum racemosum* Siebold & Zucc., which both display P/O of around 3000, while the former produced > 350 times more pollen (i.e., 2,494,654) and ovules (i.e., 704) than the latter (i.e., 6,435 pollen and 2 ovules) (Table S1). This difference is even more extreme for species with P/O around 300, as pollen production differs 3000-fold between *Papaver somniferum* and *Fumaria muralis* Sond. ex W.D.J.Koch (Table S1).

**Figure 2.**
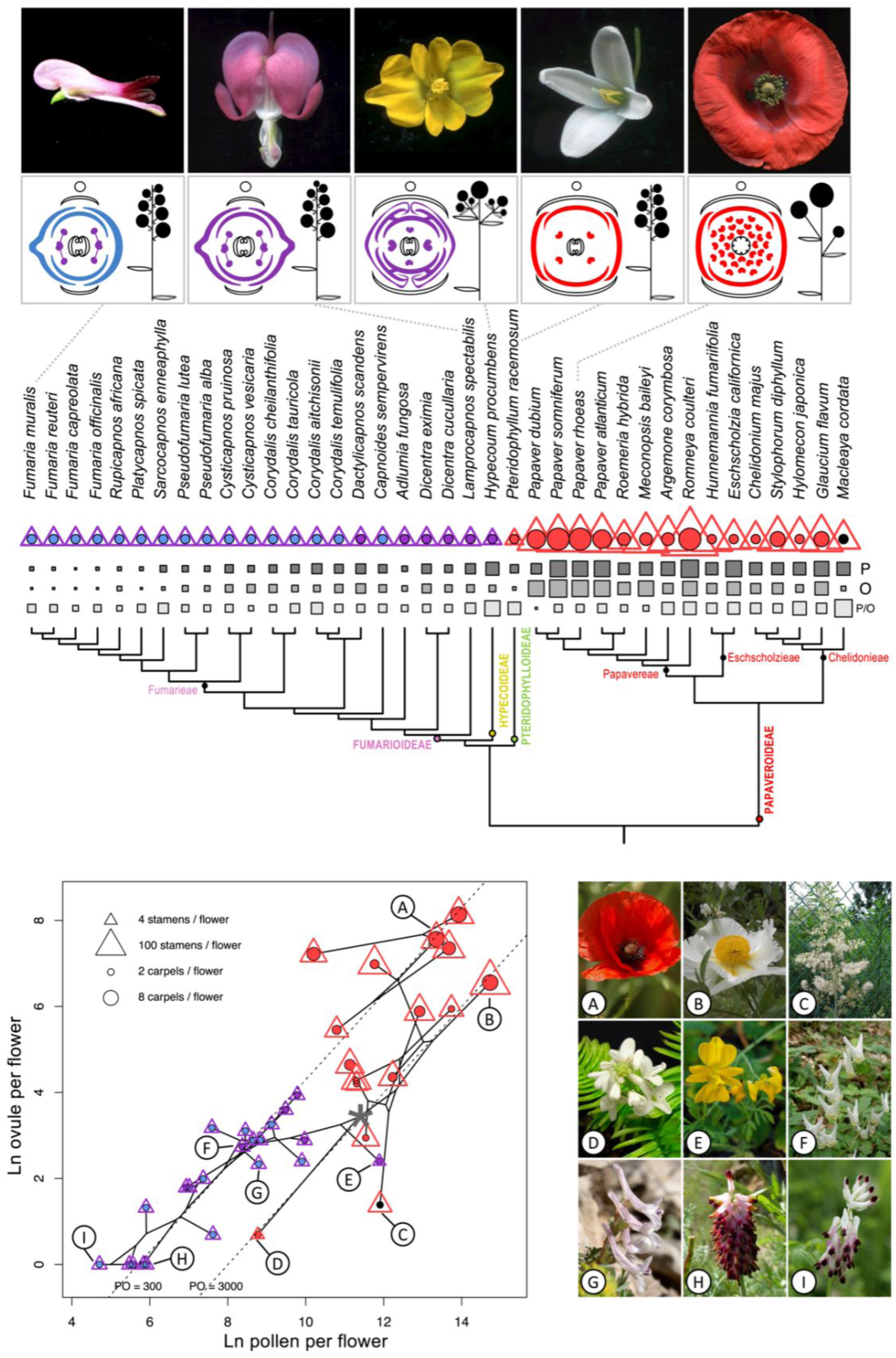
Evolutionary diversification of floral structure and symmetry in 38 Papaveraceae species. The size of each triangle is log-proportional to the stamen number per flower while the size of the circles is linearly proportional to carpel number. The colours of each reflect the symmetry of the androecium and corolla, respectively: red, actinomorphic; purple, disymmetric; blue, zygomorphic; black, apetalous. **Upper panel**: Floral traits (including pollen number per flower (P), ovule number per flower (O) and P/O ratio, with the size of the squares proportional to P, O and P/O) mapped onto the phylogeny of Papaveraceae, with pictures and floral diagrams for selected species. **Lower panel**: Phylomorphospace plot indicating relations of phylogenetically linked, ln-transformed means of pollen number per flower and ovule number per flower. Locations of internal nodes reflect ancestral states inferred based on Brownian motion. Dashed lines indicate P/O ratios of 300 and 3000. The inferred root state is marked with *. A: *Papaver rhoeas* L., B: *Romneya coulteri* Harv., C: *Macleaya cordata* R.Br., D: *Pteridophyllum racemosum* Siebold & Zucc., E: *Hypecoum procumbens* L., F: *Dicentra cucullaria* Bernh., G: *Corydalis tauricola* (Cullen & Davis) Lidén, H: *Platycapnos spicata* (L.) Bernh., I: *Fumaria capreolata* L. Photographs A: Pere Barnola; B, C, G: Wikimedia Commons, by, respectively, Lalupa (public domain), Sphl (licensed under GNU FDL), Zeynel Cebeci (licensed under CC BY-SA 4.0); D-F, H: Oriane Hidalgo; I: Jean-Marie Martin.

Pollen number, ovule number, and P/O per flower showed different evolutionary trait-associations across Papaveraceae (Fig. 3). All these traits had very high values for Pagel’s λ (Fig 3, diagonals), indicating strong phylogenetic signal in all traits when assuming Brownian Motion, enabling ancestral character state reconstruction. P/O had a stronger positive correlation with pollen than ovule number per flower (*ρ* = 0.46 vs. *ρ =* 0.30, respectively, Fig. 3). Similarly, flower number had a stronger negative correlation with ovule than pollen number per flower (*ρ* = 0.50 vs *ρ =* 0.27, respectively, Fig. 3). Both pollen and ovule number per flower correlated positively with the number of their corresponding floral organs (i.e., stamens and carpels; *ρ* = 0.59 and *ρ =* 0.63, respectively).

**Figure 3.**
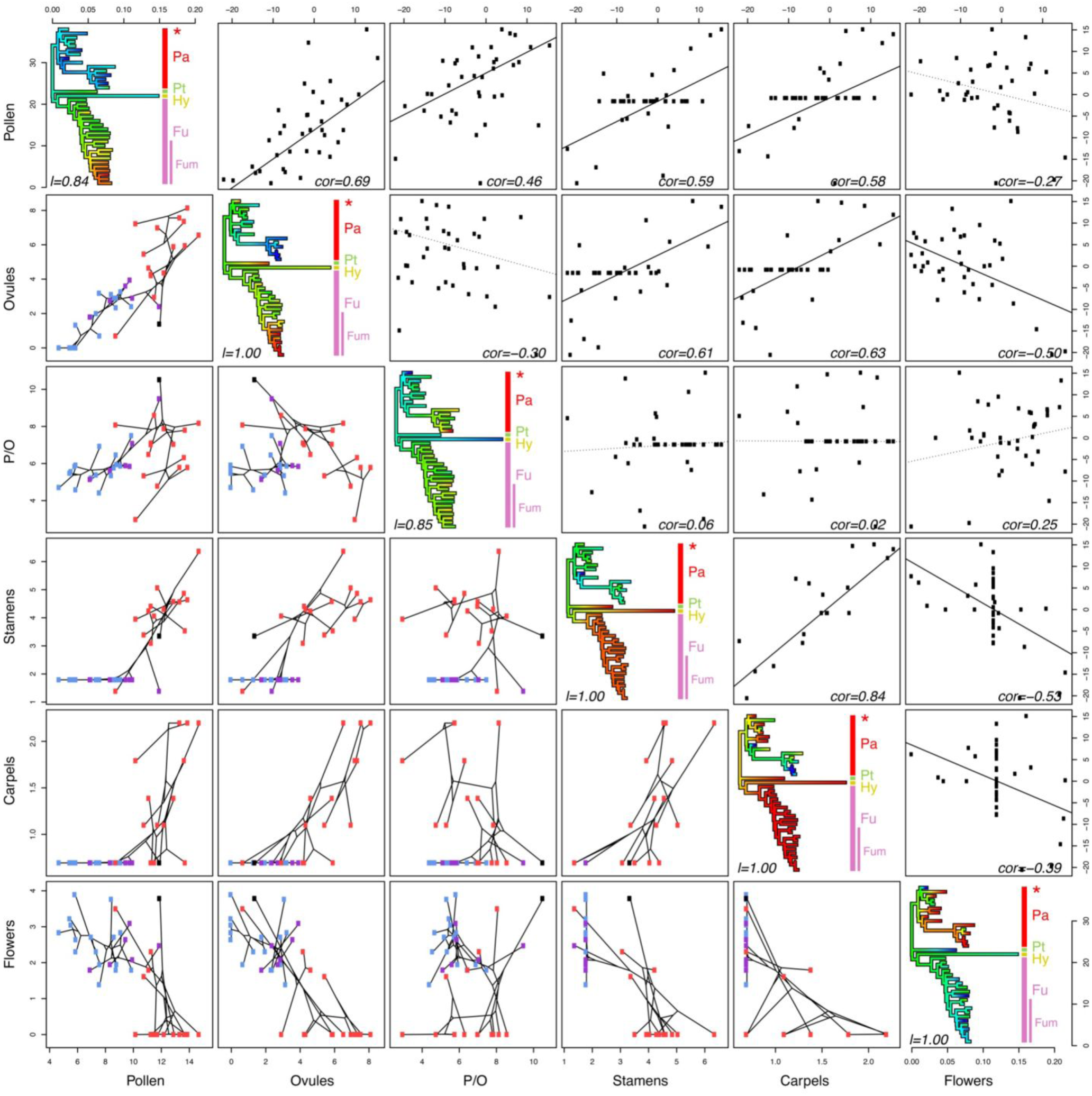
Multidimensional phylogenetic scatterplot matrix depicting phylogenetic patterns of joint diversification of six reproductive traits of Papaveraceae. Traits depicted in each panel are identified by the labels on the x and y axis in the plot margin. Diagonal elements indicate ancestral state character reconstructions of log transformed variables with values scaled linearly from red (low values) to blue (high values, given in Table 1); maximum likelihood values of lambda (λ) are indicated. Off-diagonal panels of the bottom triangle represent bivariate phylomorphospace plots with ancestral character states as in the plots on the diagonal. Tips are coloured by floral symmetry: blue, zygomorphic; purple, disymmetric; red, actinomorphic; black, apetalous. Off-diagonal panels of the upper triangle represent bivariate character correlations (with Pearson correlation coefficient indicated) of standardised phylogenetic independent contrasts inferred from the lambda-transformed trees of the diagonal panels. For visualisation purposes, an ordinary least squares regression line through the origin is drawn solid for significant (OLS, P < 0.05) associations and dotted otherwise. Clades are labelled as Pa for Papaveroideae (the asterisk indicates the position of *Macleaya*), Pt for Pteridophylloideae, Hy for Hypecoideae, Fu for Fumarioideae and Fum for Fumarieae.

**Table 1.**
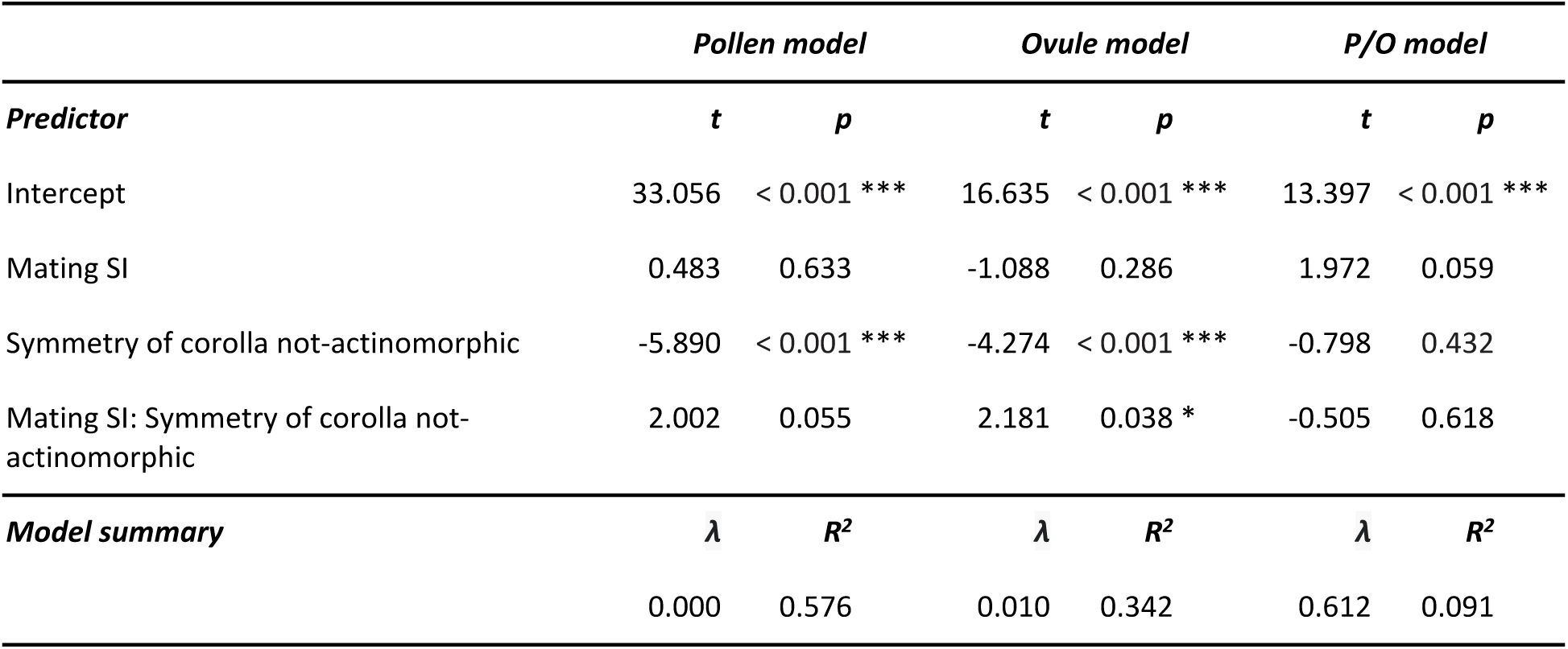
Phylogenetic least-squares regressions examining the effect of mating system and floral symmetry and their interaction on pollen number (P), ovule number (O) and their P/O ratio. SI: self-incompatibility. (* = significant at the α < 0.05 level; ** = significant at the α < 0.01 level; *** = significant at the a α < 0.001 level).

Pollen and ovule production per flower was significantly lower in non-actinomorphic species compared to actinomorphic species (Table 1; Fig. 4). Self-incompatibility was associated with a higher ovule production per flower in non-actinomorphic species, but not in actinomorphic ones (Fig. 4; Table 2). Importantly, pollen and ovule numbers displayed clear and contrasting evolutionary trends between actinomorphic and non-actinomorphic species, but these patterns were masked when analysing only their P/O ratio, evidenced by a low R-squared of 0.091 for the P/O model, compared to 0.576 and 0.342 for the pollen and ovule models, respectively (Table 1).

**Figure 4.**
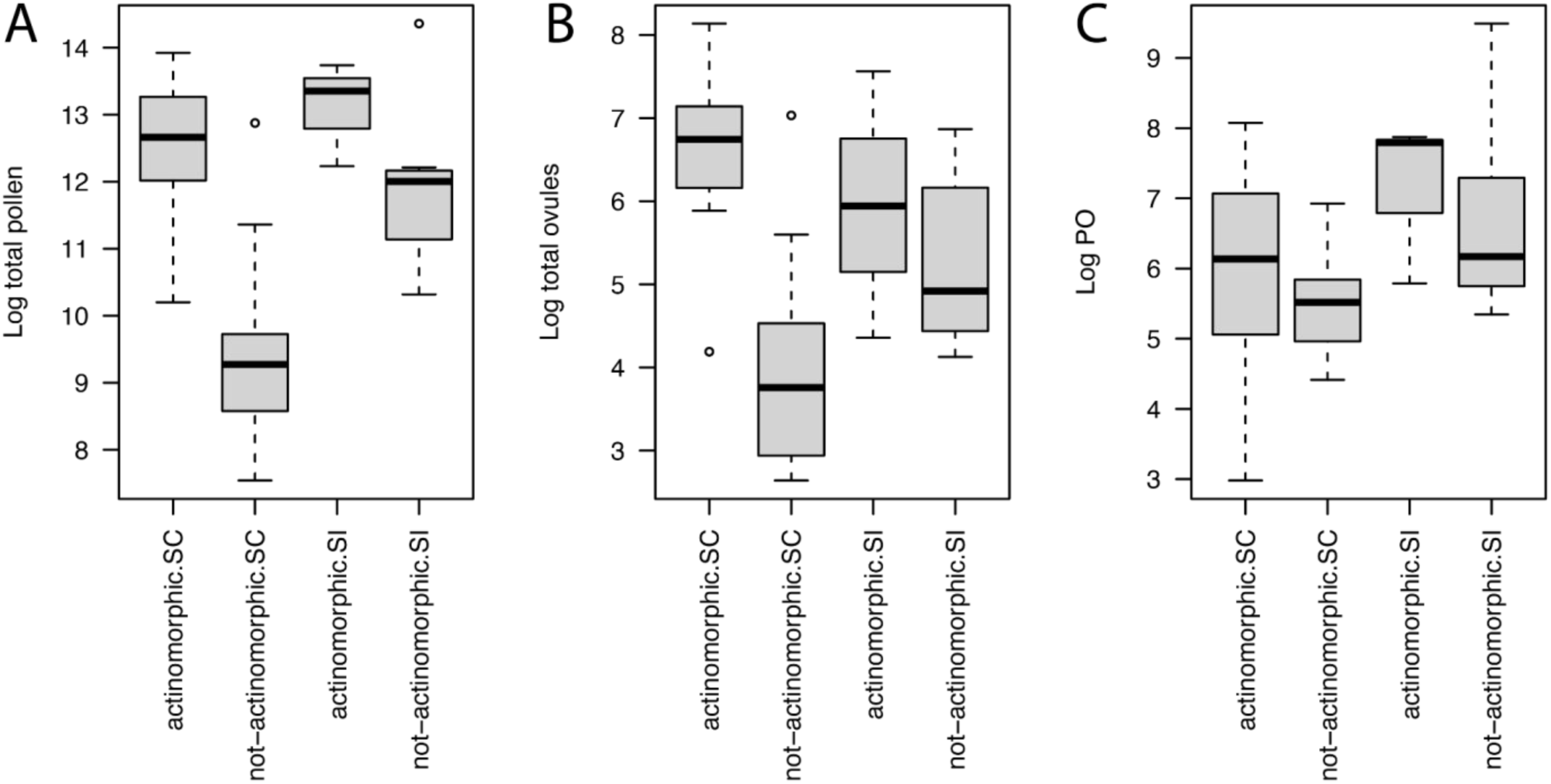
Comparisons of the mean values for (A) the number of pollen grains per flower (B) the number of ovules per flower and (C) P/O ratios for the 38 Papaveraceae species analysed, split by floral symmetry (actinomorphic or not) and self-compatibility.

**Table 2.** Average values for pollen and ovule numbers, P/O ratio, and number of stamens (ST), carpels (CA) and flowers per inflorescence (Flo) in Papaveraceae taxa (raw data are provided in Table S1); standard deviations are indicated following the ± sign, rounded off to zero decimals. The order of clades follows the evolutionary history obtained with the rooted phylogeny used for this study (Figure 3). Values for genera with multiple species were calculated with the average values for species within that genus. Values for tribes, subfamilies and the family were calculated using average values for genera. The number of flowers used for calculating averages are indicated for each species and genus. For pollen numbers, only measurements obtained with flow cytometry were considered. ^a^Self-compatibility/incompatibility is indicated for species and genera where this is known (references in Table S1). *: Multiple accessions analysed.

| Taxa | Pollen | Ovules | P/O | ST | CA | Flo | Inc<sup>a</sup> |
| --- | --- | --- | --- | --- | --- | --- | --- |
| Papaveraceae | 219,350 | 206 | 2,990 | 54 | 3 | 11 | - |
| Papaveroideae | 456,401 | 431 | 4,669 | 111 | 4 | 6 | - |
| Chelidoniae | 161,933 | 111 | 9,098 | 55 | 3 | 12 | - |
| *Macleaya* (N=4) | 148,133 ± 26,845 | 4 | 37,033 ± 6,711 | 29 ± 5 | 2 | 44 | ? |
| *Glaucium* (N=3) | 409,093 ± 63,114 | 360 ± 30 | 1,140 ± 168 | 97 ± 11 | 4 | 1 | SC |
| *Hylomecon* (N=1) | 102,660 | 19 | 5,403 | 58 | 2 | 1 | ? |
| *Stylophorum* (N=3) | 68,323 ± 9,427 | 104 ± 13 | 679 ± 180 | 69 ± 4 | 4 | 6 | SC |
| *Chelidonium* (N=4) | 81,457 ± 12,281 | 67 ± 5 | 1,235 ± 255 | 22 ± 2 | 2 | 10 | SC |
| Eschscholziae | 502,739 | 227 | 1,862 | 58 | 2 | 1 | - |
| *Eschscholzia* (N=4) | 924,999 ± 197,864 | 381 ± 104 | 2,605 ± 758 | 34 ± 2 | 2 | 1 | SI |
| *Hunnemannia* (N=1) | 80,478 | 72 | 1,118 | 82 | 2 | 1 | ? |
| Papaverae | 634,526 | 833 | 1,363 | 188 | 5 | 2 | - |
| *Romneya* (N=1) | 2,494,654 | 704 | 3,544 | 581 | 9 | 1 | ? |
| *Argemone* (N=1) | 205,028 | 78 | 2,629 | 74 | 3 | 1 | SI |
| *Meconopsis* (N=3) | 128,783 ± 69,961 | 1,078 ± 389 | 111 ± 38 | 158 ± 38 | 3 | 1 | SC |
| *Roemeria* (N=3) | 48,698 ± 9,435 | 234 ± 81 | 226 ± 72 | 30 ± 4 | 3 | 5 | SC |
| *Papaver* (4 spp.) | 784,503 | 2,069 | 307 | 96 | 8 | 1 | SC |
| *P. atlanticum* (N=1) | 867,717 | 1,560 | 556 | 130 | 6 | 1 | ? |
| *P. dubium* (N=1) | 26,918 | 1,368 | 20 | 52 | 6 | 1 | SC |
| *P. rhoeas* (N=2) | 628,742 ± 78,483 | 1,926 ± 126 | 326 ± 20 | 96 ± 4 | 9 | 1 | SI |
| *P. somniferum* (N=3) | 1,113,133 ± 125,997 | 3,420 ± 135 | 326 ± 39 | 104 ± 14 | 9 ± 1 | 1 | SC |
| *Pteridophyllum* (N=4) | 6,435 ± 1,349 | 2 | 3,218 ± 675 | 4 | 2 | 33 | SC |
| *Hypecoum* (N=6*) | 145,488 ± 29,972 | 11 ± 2 | 13,223 ± 1,904 | 4 | 2 | 10 ± 4 | SI |
| Fumarioideae | 6,198 | 15 | 439 | 6 | 2 | 15 | - |
| **Lamprocapnos** (N=3) | 21,415 ± 5,774 | 18 | 1,190 ± 321 | 6 | 2 | 8 | SI |
| **Dicentra** (2 spp.) | 10,193 | 27 | 322 | 6 | 2 | 11 | SI |
| *D. cucullaria* (N=2) | 4,331 ± 364 | 16 ± 1 | 281 ± 33 | 6 | 2 | 7 | SI |
| *D. eximia* (N=4) | 13,125 ± 453 | 37 ± 3 | 362 ± 29 | 6 | 2 | 14 | SI |
| **Adlumia** (N=2) | 1,026 ± 14 | 6 | 171 ± 2 | 6 | 2 | 6 | ? |
| **Capnoides** (N=4) | 6,684 ± 2614 | 18 ± 3 | 414 ± 187 | 6 | 2 | 7 | ? |
| **Dactylicapnos** (N=4) | 17,757 ± 8,265 | 52 ± 3 | 338 ± 136 | 6 | 2 | 22 | SC |
| **Corydalis** (4 spp.) | 9,090 | 17 | 742 | 6 | 2 | 18 | SI |
| *C. temulifolia* (N=4) | 9,171 ± 363 | 26 ± 3 | 361 ± 60 | 6 | 2 | 17 | SI |
| *C. aitchisonii* (N=2) | 19,301 ± 2,934 | 11 | 1,755 ± 267 | 6 | 2 | 6 | SI |
| *C. tauricola* (N=3) | 6,580 ± 1,066 | 10 | 640 ± 119 | 6 | 2 | 6 | SI |
| *C. cheilanthifolia* (N=3) | 4,685 ± 215 | 22 ± 3 | 212 ± 22 | 6 | 2 | 43 | SI |
| **Fumariae** | 1,368 | 6 | 349 | 6 | 2 | 18 | - |
| **Cysticapnos** (2 spp.) | 3,853 | 21 | 219 | 6 | 2 | 10 | SC |
| *C. vesicaria* (N=2) | 1,984 ± 1,750 | 24 ± 8 | 120 ± 113 | 6 | 2 | 4 | SC |
| *C. pruinosa* (N=2) | 5,722 ± 108 | 18 | 318 ± 6 | 6 | 2 | 15 | SC |
| **Pseudofumaria** (2 spp.) | 1,397 | 7 | 201 | 6 | 2 | 13 | SC |
| *P. alba* (N=2) | 1,110 ± 121 | 7 | 185 ± 20 | 6 | 2 | 15 | SC |
| *P. lutea* (N=3) | 1,589 ± 30 | 7 ± 1 | 217 ± 11 | 6 | 2 | 10 | SC |
| **Sarcocapnos** (N=3) | 2,036 ± 159 | 2 | 1,018 ± 80 | 6 | 2 | 7 | SC |
| **Platycapnos** (N=2) | 344 ± 115 | 1 | 344 ± 115 | 6 | 2 | 49 | SC |
| **Rupicapnos** (N=4) | 367 ± 209 | 4 | 99 ± 54 | 6 | 2 | 10 | SC |
| **Fumaria** (4 spp.) | 213 | 1 | 213 | 6 | 2 | 19 | SC |
| *F. officinalis* (N=3) | 239 ± 159 | 1 | 239 ± 159 | 6 | 2 | 25 | SC |
| *F. capreolata* (N=4) | 64 ± 11 | 1 | 64 ± 11 | 6 | 2 | 17 | SC |
| *F. reuteri* (N=1) | 260 | 1 | 260 | 6 | 2 | 21 | SC |
| *F. muralis* (N=3) | 369 ± 159 | 1 | 369 ± 159 | 6 | 2 | 14 | SC |

## Discussion

Our study is based on our new protocol for pollen counting using flow cytometry, considerably improving the accuracy of pollen counts (Fig. 1) and enabling a higher throughput compared to manual counts as performed in previous assessments. Given the wide availability of flow cytometers in many institutions (compared to, e.g., Coulter counters), we expect that our method will be broadly applicable in many laboratories.

We highlight that our survey of Papaveraceae revealed a much broader variation not only in pollen production, but also in ovule production and P/O ratios, than previously reported in studies that focused on only a few species (e.g., Cruden, 1977; Hannan, 1981; Dahl, 1989; Ohara & Higashi, 1994; Fukuhara, 2000; Salinas & Suárez, 2003; Erbar & Langlotz, 2005; Xiao *et al*., 2016; Wu *et al*., 2019). Nevertheless, despite their limited taxon sampling, these previous studies did report particularly large numbers of pollen grains per flower in some representatives of the Papaveraceae subfamily Papaveroideae (e.g., ca. 450,000 to over two millions in *Platystemon californicus* Benth., Hannan, 1981; ca. 2,636,000 in *Papaver rhoeas*, Erbar & Langlotz, 2005), amongst the highest known for insect-pollinated flowers (Erbar & Langlotz, 2005). In agreement with these results, we found similarly extreme values in the Papaveroideae, though not in the same species, with pollen counts of > 2 million per flower for *Romneya coulteri* and > 1 million for *Eschscholzia californica* Cham. and *Papaver somniferum* (Table S1). Our counts for *Papaver rhoeas*, on the other hand, represented a fourth of the previously reported values at c. 600,000 (Table 2). This difference may be due to intraspecific variation in the number of pollen, as indicated above for *Platystemon californicus*, and also observed in our dataset (some taxa show an intraspecific variation of a similar magnitude, and even largely exceeding 4x, see Table S1). It could also result from a tendency to overestimate pollen numbers with manual counting protocols (Figure 2D). The lowest pollen counts (< 100 per flower; Tables S1-S3) were obtained in *Fumaria capreolata*, a taxon that had not been studied in this respect so far, thus considerably extending the range of variation known for Papaveraceae (Fig. 2; Table S1). Importantly, we demonstrate that interspecific variation in absolute pollen and ovule production significantly exceeds that of their ratio, P/O. This underlines that the evolution of pollen and ovule production is multifarious, more so than the patterns observed by analysing just P/O would suggest. This aligns with the findings of Harder and Johnson (2023) regarding the limited usefulness of P/O as an indicator of the mating system. Our results reveal that the great variation in pollen and ovule production (as well as P/O) shows contrasting trait-associations, highlighting distinct evolutionary trajectories between and within subfamilies (Figs. 2 & 3). Multiple patterns of evolutionary trait associations arise in different clades of the family (Fig. 5), which we highlight below through five specific examples. The first two examples highlight contrasting underlying pollen and ovule trajectories resulting in shifts or stasis in the P/O ratio, while the latter three highlight contrasting trait associations of pollen, ovules and P/O ratios.

**Figure 5.**
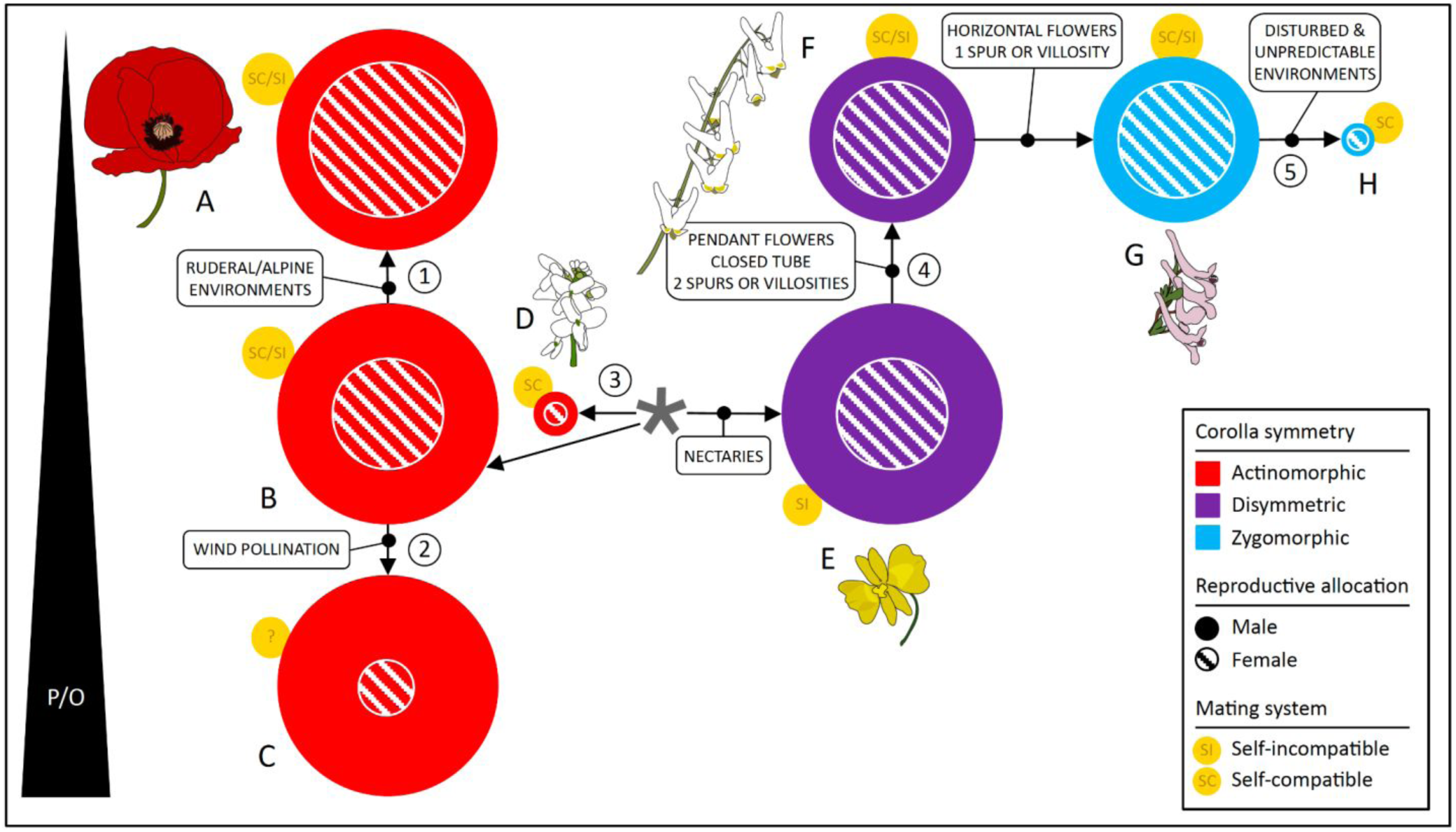
Schematic representation of transitions in reproductive allocation, floral symmetry and mating system in the Papaveraceae. The inferred ancestral state is marked with *. A: *Papaver rhoeas* L., B: *Romneya coulteri* Harv., C: *Macleaya cordata* R.Br., D: *Pteridophyllum racemosum* Siebold & Zucc., E: *Hypecoum procumbens* L., F: *Dicentra cucullaria* Bernh., G: *Corydalis tauricola* (Cullen & Davis) Lidén, H: *Platycapnos spicata* L. (see pictures in Figure 2). (For details of the different shifts in trait values indicated by numbers see the Discussion).

### An increased ovule production led to a drop in P/O in a Papaveroideae clade which includes alpine Meconopsis Vig. and ruderal Papaver L. and Roemeria Medik. (see Fig. 2; and shift 1 of Fig. 5)

This trend fits the expectations of a strategy for species with small seeds (e.g., ≤ 1 mm ø in *Papaver rhoeas*), whereby ovule –and hence the potential for seed– production is maximised, enabling seed bank establishment and optimising seed set returns from erratic pollinator visits (e.g., as reported in *Mimulus* L., Phrymaceae; Ritland & Ritland, 1989). In the context of ruderal or high elevation environments where pollinator visits are unpredictable, producing densely packed ovaries and many small seeds reduces the need for numerous pollinator visits and promotes a high seed set (Burd *et al*., 2009). Moreover, the likelihood of seedling establishment is likely very low due to patchy availability of suitable habitat, further promoting the advantage of having a high seed set (Körner, 2021). Nevertheless, as a larger number of offspring arise from the same parents, genetic diversity may be negatively affected, a fact further accentuated by only a moderate pollen and seed dispersal ability of these poppies (Miller *et al*., 2005). In self-incompatible species, increasing relatedness would lower the probability that a pollinator brings a compatible pollen (e.g., in *Papaver rhoeas*; Brooks, Tobias & Lawrence, 1996), and this may limit seed production in species with mixed mating (e.g., in *Papaver dubium*; Humphreys & Gale, 1974). Increased ovule production as observed in these species could then compensate for such effects.

### A drastic reduction in both pollen and ovule production resulted in no change in P/O in flowers of Pteridophyllum racemosum (see Fig. 2; and shift 3 of Fig. 5)

The genus *Pteridophyllum* Siebold & Zucc., whose phylogenetic affinities with subfamilies Papaveroideae and Fumarioideae remain unclear (Hoot *et al*., 1997; Hoot, Wefferling & Wulff, 2015; Wang *et al*., 2009; Pérez-Gutiérrez *et al*., 2012, 2015a; Sauquet *et al*., 2015), has flowers with low pollen and ovule numbers (Table 2) as observed in Fumarioideae, but a high P/O as in the Papaveraceae backbone (Fig. 2). *Pteridophyllum* is a self-compatible genus (Ozawa, Kudoh & Kachi, 2001) with oligandrous (i.e., < 10 stamens) actinomorphic flowers, that has the peculiarity of displaying a racemose inflorescence –otherwise exclusively found in Fumarioideae (Hidalgo & Gleissberg, 2010). The (re)distribution of reproductive allocation throughout the inflorescence, also observed in *Macleaya* R.Br. and some Fumarioideae [e.g., *Platycapnos spicata* (L.) Bernh.], suggests a parallel evolution of this trait, that is independent from the floral symmetry and P/O contexts.

### Wind-pollination is associated with the predicted increase in pollen production at the inflorescence level, but not at the flower level in Macleaya cordata (see Fig. 2; and shift 2 of Fig. 5)

Wind-pollinated species rely upon dispersal of large amounts of pollen, which was long thought to be related to the relatively untargeted method of pollen transfer to the carpel (Friedman & Barrett, 2009), and resulting in a reduction in the number of ovules per flower (Friedman & Barrett, 2011). In *Macleaya cordata*, a shift to wind-pollination is accompanied with a decrease in ovule number but not in pollen production per flower (Fig. 2). The estimated high P/O (Cruden, 2000; Culley, Weller & Sakai, 2002) for this species –with an average of 37,033, the highest in Papaveraceae (Table 2)– is the result of strongly decreased female allocation rather than increased male allocation. Reduced pollen production per flower in *Macleaya cordata* was however counterbalanced by a dramatic increase in the number of flowers, organised in a diffuse panicle suited for wind-pollination (Friedman & Harder, 2004; Harder & Prusinkiewicz, 2013). Indeed, the average pollen production per inflorescence in the species was estimated to exceed six million grains (see Table S1 for the number of flowers per inflorescence and pollen count per flower), more than any other inflorescence or solitary flower in the family examined to date. In addition, a shift to wind pollination is also reflected in pollen morphology, as *Macleaya cordata* has been reported to have the smallest pollen in the tribe Chelidonieae, and a finger-like structure around the pore interpreted as an adaptation to wind-pollination that could enhance harmomegathy (i.e. the ability of pollen to absorb bending stresses such as occurring during desiccation and rehydration; Muller, 1979) (Suárez-Santiago et al., 2018). Reduced pollen size in this species is also consistent –here at inflorescence level– with the documented trade-off between the number and size of pollen grains, as it is associated with the production of high pollen numbers (Cruden & Miller-Ward, 1981; Vonhof & Harder, 1995).

### Transition to disymmetric flowers is not *per se* associated with a reduced P/O, but only when accompanied with pollinator type-restricting traits (Fig. 2; shift 4 of Fig. 5)

*Hypecoum procumbens* L., sister to Fumarioideae, presents pollen and ovule numbers similar to the ancestral values reconstructed for Papaveraceae (Fig. 2), including a high P/O ratio, despite it displaying the derived trait of possessing disymmetric flowers. This pattern suggests that evolution of disymmetry did not impact pollen and ovule production by itself. Floral orientation in *Hypecoum* L. is similar to Papaveroideae, with erect flowers, open corolla and freely accessible anthers, which can attract a range of pollinator types. In stark contrast, disymmetric Fumarioideae display a drop in P/O, due to reduced pollen production likely associated with traits that restricted the morphological and behavioural range of insect visitors. These are in particular the change in floral orientation from erect to pendant (*Ehrendorferia* Fukuhara & Lidén excepted; Fenster, Armbruster & Dudash, 2009) and the evolution of a closed corolla and spurs (Woźniak & Sicard, 2018). *Hypecoum* also differs from Fumarioideae in pollen morphology, e.g., in the exine ornamentation and aperture number (Pérez-Gutiérrez *et al*., 2015b), suggesting a profound developmental remodelling that impacted the entire reproductive system.

### Transition to zygomorphy has no noticeable impact on P/O, however, a drastic reduction in ovule and pollen number in the core Fumarieae is associated with the unique combination of zygomorphy with self-compatibility and adaptation to disturbed and unpredictable environments (Fig. 2; shift 5 of Fig. 5)

The shift from disymmetry to zygomorphy arose with the reorientation of the inflorescence from horizontal to vertical, and of the flowers, from pendant to horizontal (e.g., *Lamprocapnos* Endl. vs. *Corydalis*). The derived configuration is thought to enhance recognition of inflorescences by hymenopteran pollinators, ensure consistent directionality of pollinator movement on the inflorescence (Harder *et al*., 2004), and improve precision of pollen placement on pollinators (Wang *et al*., 2014). Surprisingly, for most species in Fumarieae, the transition from disymmetry to zygomorphy has no noticeable impact on pollen and ovule production, even when associated with self-compatibility as seen across Fumarieae (all Fumarieae are self-compatible; Lidén, 1986). A drastic reduction in both pollen and ovule number is, however, observed in a clade of Fumarieae containing *Fumaria*, *Platycapnos*, *Rupicapnos* Pomel, and *Sarcocapnos* DC. (Fig. 2). These small-flowered taxa are adapted to life in disturbed and unpredictable environments and each presents an extremely low pollen and ovule number per flower (e.g., *Fumaria*, Fig. 2), possibly displaying a “selfing syndrome” (e.g., Ritland & Ritland, 1989; Sicard & Lenhard, 2011; de Vos *et al*., 2014b). In this context, self-compatibility may enable such reduced reproductive investment, rather than necessarily inducing it.

## Concluding remarks

We have presented the first family-wide quantitative assessment of male reproductive investment in the Papaveraceae, which together with data on additional floral traits, have highlighted very distinctive evolutionary trajectories both between and within subfamilies (Fig. 5). This was enabled by developing an accessible, accurate, and high-throughput method for assessing male reproductive investment in plants. The method has been validated by comparisons with manual counts and has been shown to considerably reduce the standard error in pollen counts (Fig. 2E), thereby preventing overestimations of pollen numbers. Furthermore, the method was shown to be rapid, with the possibility of analysing > 50 pollen samples in one day. Such high throughput represents a significant increase in time-efficiency for estimating male investment in reproduction, allowing us to substantially increase the sample size in our study. Although we recognise that our taxonomic sampling is still incomplete, the importance of our considerably broader survey compared to previous studies for understanding the ecology and evolution of plant reproductive diversity is illustrated by five examples of interactions among multiple floral traits during evolution. Thus, our study demonstrates the impact such high-throughput family-wide assessments of male reproductive investment can have. We therefore expect our protocol to be of use for the wider community of pollination ecologists and evolutionary biologists studying male reproductive investment in plants.

## Acknowledgements

We thank the following funding sources: Emily Holmes Memorial Foundation scholarship (Y.W.); the Harding Alpine Plant Conservation & Research Programme through Winton Philanthropies and Swiss National Science Foundation grant n°310030_185251 (J.M.d.V.); and the ‘Ajuts a Grups de Recerca Consolidats’ (2021SGR00315) from the Generalitat de Catalunya (O.H.). We thank the following botanical gardens for providing plant material: Royal Botanic Gardens Kew, Botanische Tuinen Universiteit Utrecht, Cambridge University Botanic Garden, Botanical Garden of Barcelona, and the collaborators who provided field assistance and sample supply, in particular Teresa Garnatje, Jordi Luque, Maarten Christenhusz, Joan Vallès and Luca Pegoraro. We thank Thais Vasconcelos for her assistance with the protocol for manual pollen grain counting.

## Author contributions

O.H., J.M.d.V. and I.J.L. designed the study and developed the pollen counting method by flow cytometry; O.H. carried out the flow cytometry pollen counts; Y.W. carried out the manual pollen counts, the floral characterisation and the phylogeny reconstruction; J.M.d.V. performed comparative analyses; all authors interpreted the data and contributed to manuscript preparation.

---

**Table S1.**
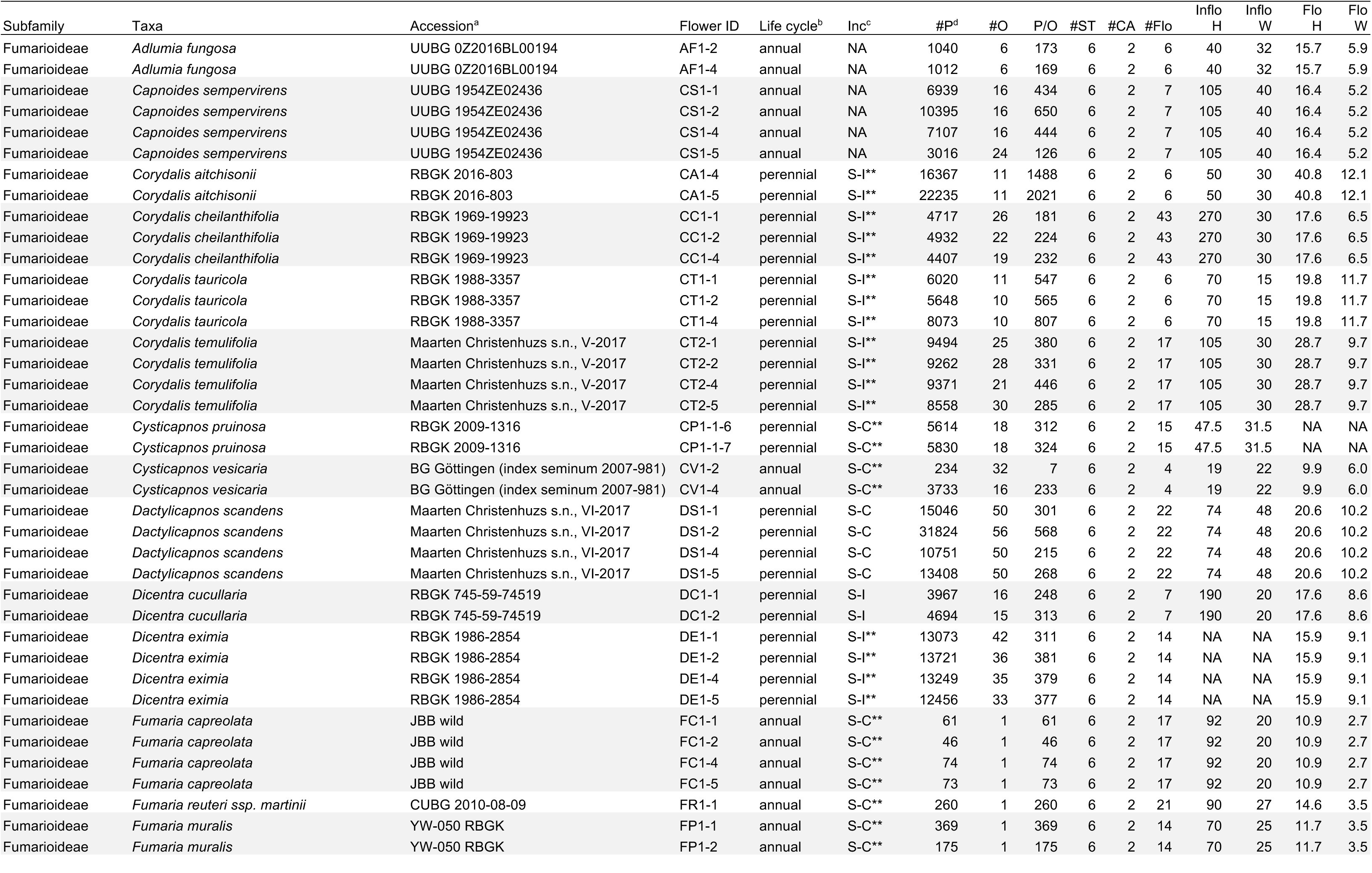
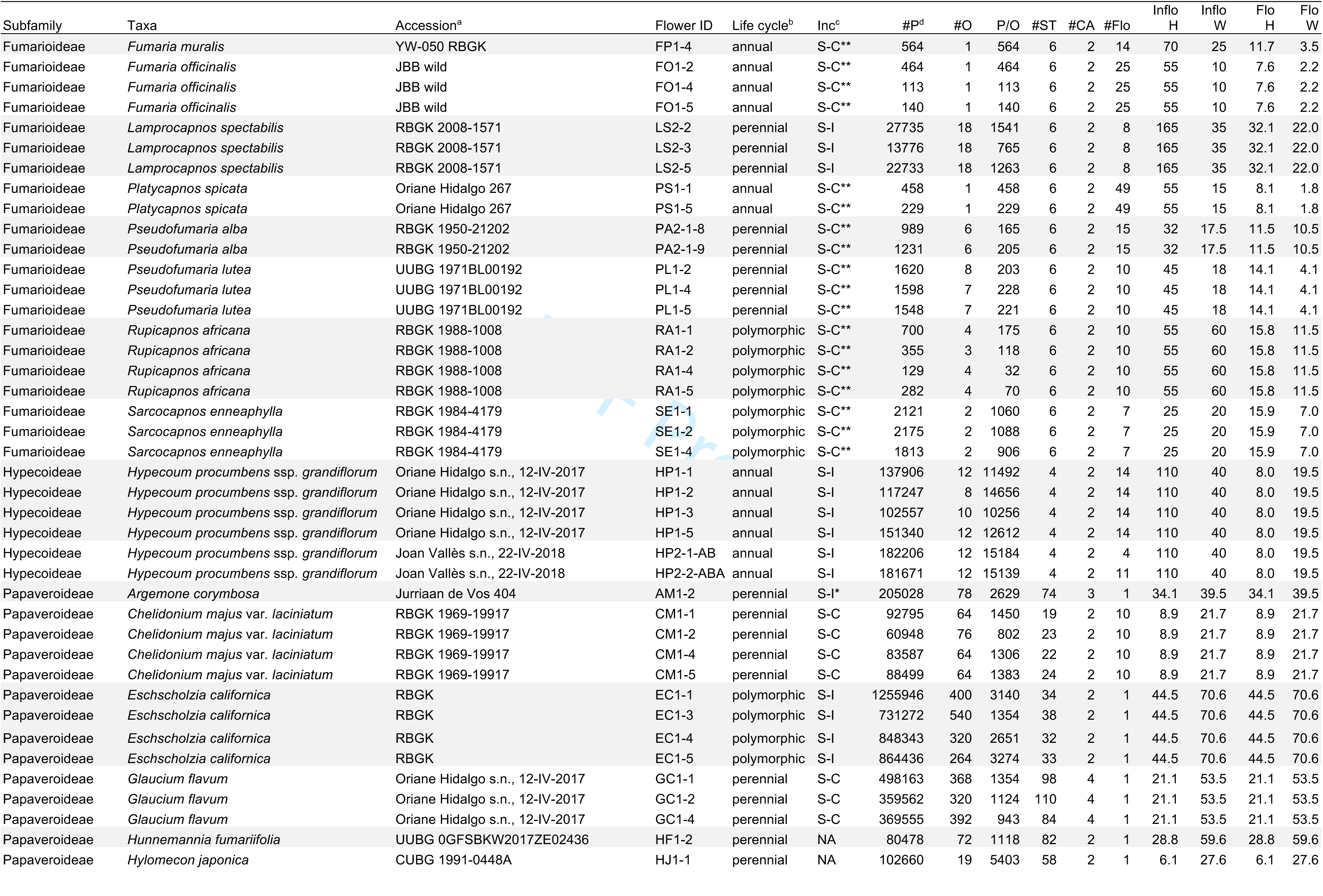
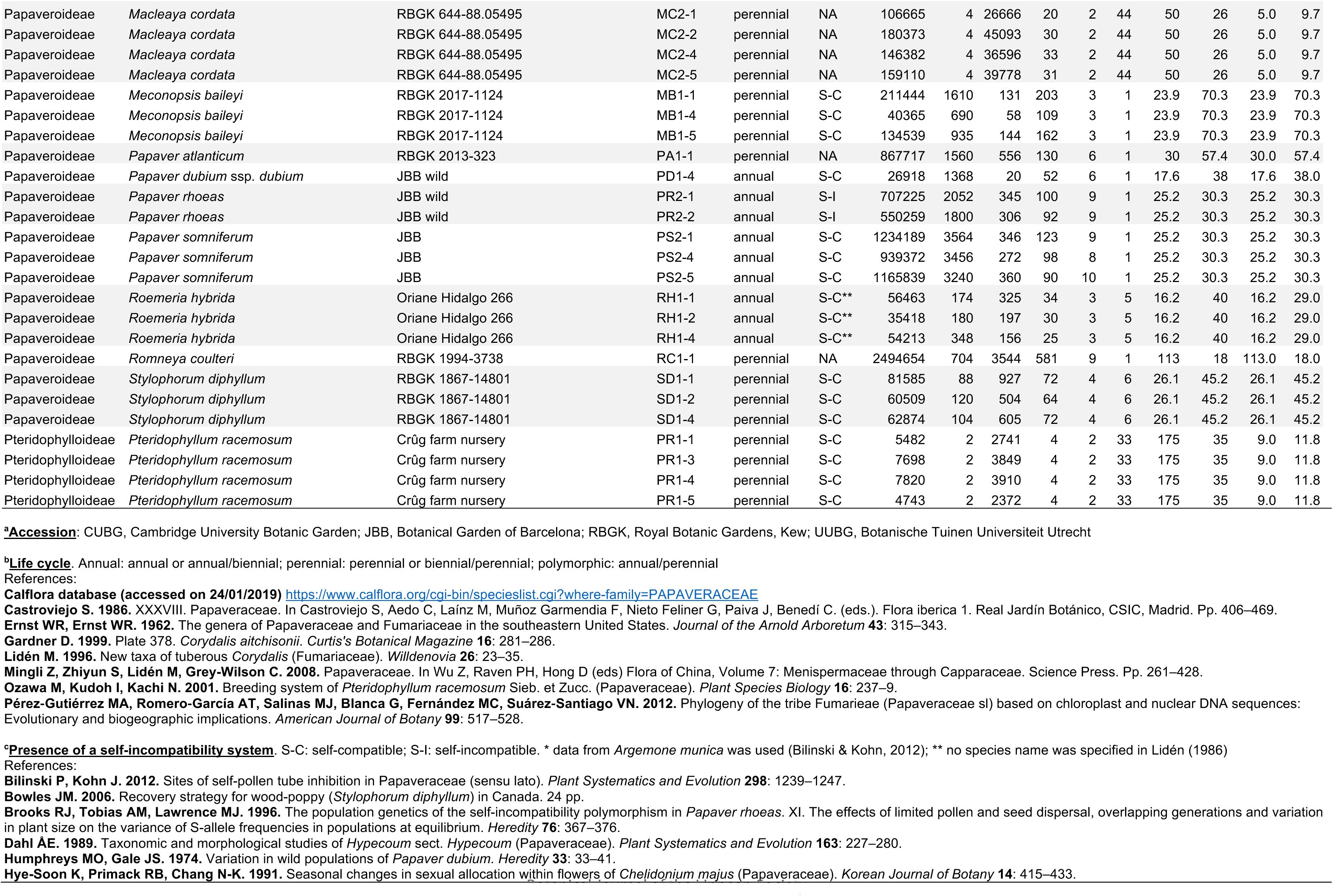
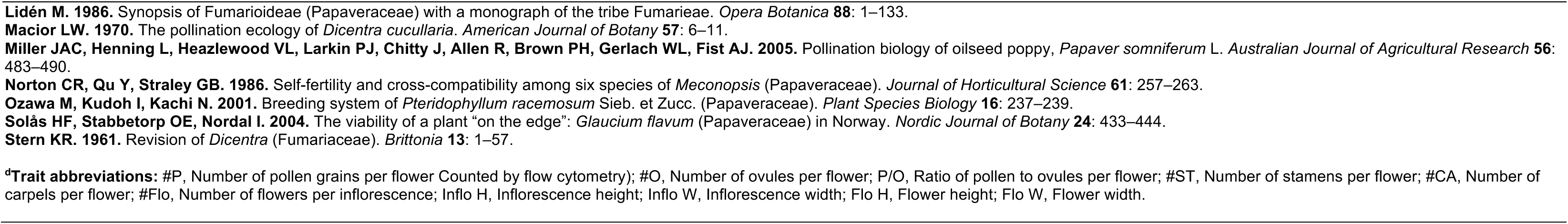
Summary of the floral and inflorescence traits and mating system for each of the 38 species with FC pollen count included in the analyses.

**Table S2.** GenBank accessions used for phylogenetic reconstruction representing the 38 species with FC pollen count and three outgroups.

| Species | matK | rbcl |
| --- | --- | --- |
| *Adlumia fungosa* |  | KM364740 |
| *Argemone mexicana* (i.o. *A. corymbosa*) | GQ434281 | U86621 |
| *Capnoides sempervirens* | KP715386 | MG249112 |
| *Chelidonium majus* | KP715387 | HM849887 |
| *Corydalis cheilanthifolia* | KP715389 |  |
| *Corydalis incisa* (i.o. *C. aitchisonii*) |  | DQ912903 |
| *Corydalis solida* (i.o. *C. tauricola*) | KP715392 |  |
| *Corydalis temulifolia* |  | KX272438 |
| *Cysticapnos pruinosa* |  | KM364747 |
| *Cysticapnos vesicaria* | LN610931 | KM364748 |
| *Dactylicapnos scandens* | LN610934 | KX527241 |
| *Dicentra cucullaria* | LN610936 | KM364751 |
| *Dicentra eximia* | DQ182345 | L37917 |
| *Eschscholzia californica* | KP715395 | HM849984 |
| *Fumaria capreolata* | KP715397 | HM850014 |
| *Fumaria reuteri* ssp. *martinii* |  | KX678375 |
| *Fumaria muralis* | HM851023 | HM850015 |
| *Fumaria officinalis* | JN895721 | JN893098 |
| *Glaucium flavum* | JN895721 | U86626 |
| *Hunnemannia fumariifolia* | LN614544 | U86627 |
| *Hylomecon japonica* | FJ626522 | FJ626611 |
| *Hypecoum procumbens* | KP715399 |  |
| *Lamprocapnos spectabilis* | KP715400 | DQ912901 |
| *Macleaya microcarpa* (i.o. *M. cordata*) | FJ626523 | FJ626612 |
| *Meconopsis quintuplinervia* (i.o. *M. baileyi*) | FJ626524 | FJ626613 |
| *Papaver dubium* subsp. *dubium* | JN893923 | HM850229 |
| *Papaver hybridum* (i.o. *P. atlanticum*) | JN895926 | JN890902 |
| *Papaver rhoeas* | KP715401 | HM850230 |
| *Papaver somniferum* | MG221004 | HM850231 |
| *Platycapnos spicata* | KP715402 |  |
| *Pseudofumaria alba* | KP715403 | KM364762 |
| *Pseudofumaria lutea* | KP715404 | KM364763 |
| *Pteridophyllum racemosum* |  | U86631 |
| *Roemeria hybrida* |  | KX364740 |
| *Romneya coulteri* |  | U86632 |
| *Rupicapnos africana* | KP715405 | LN610958 |
| *Sarcocapnos enneaphylla* | KP715406 | KM364765 |
| *Stylophorum diphyllum* | JX087859 | JX087704 |
| *Euptelea polyandra* | DQ401348 | L12645 |
| *Thalictrum javanicum* | DQ478615 | AY954496 |
| *Magnolia grandiflora* | AF548640 | AF119180 |

**Table S3.**
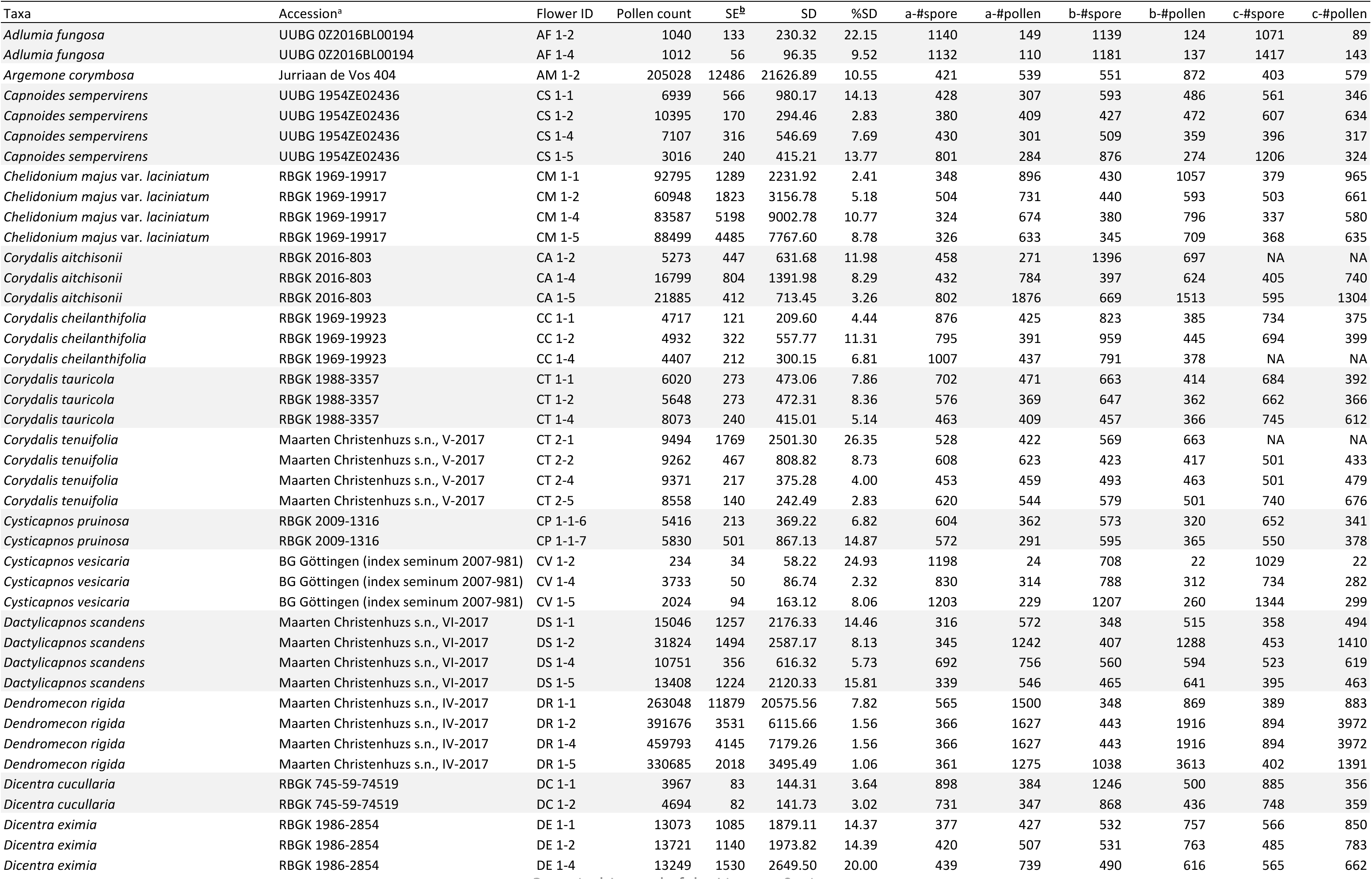
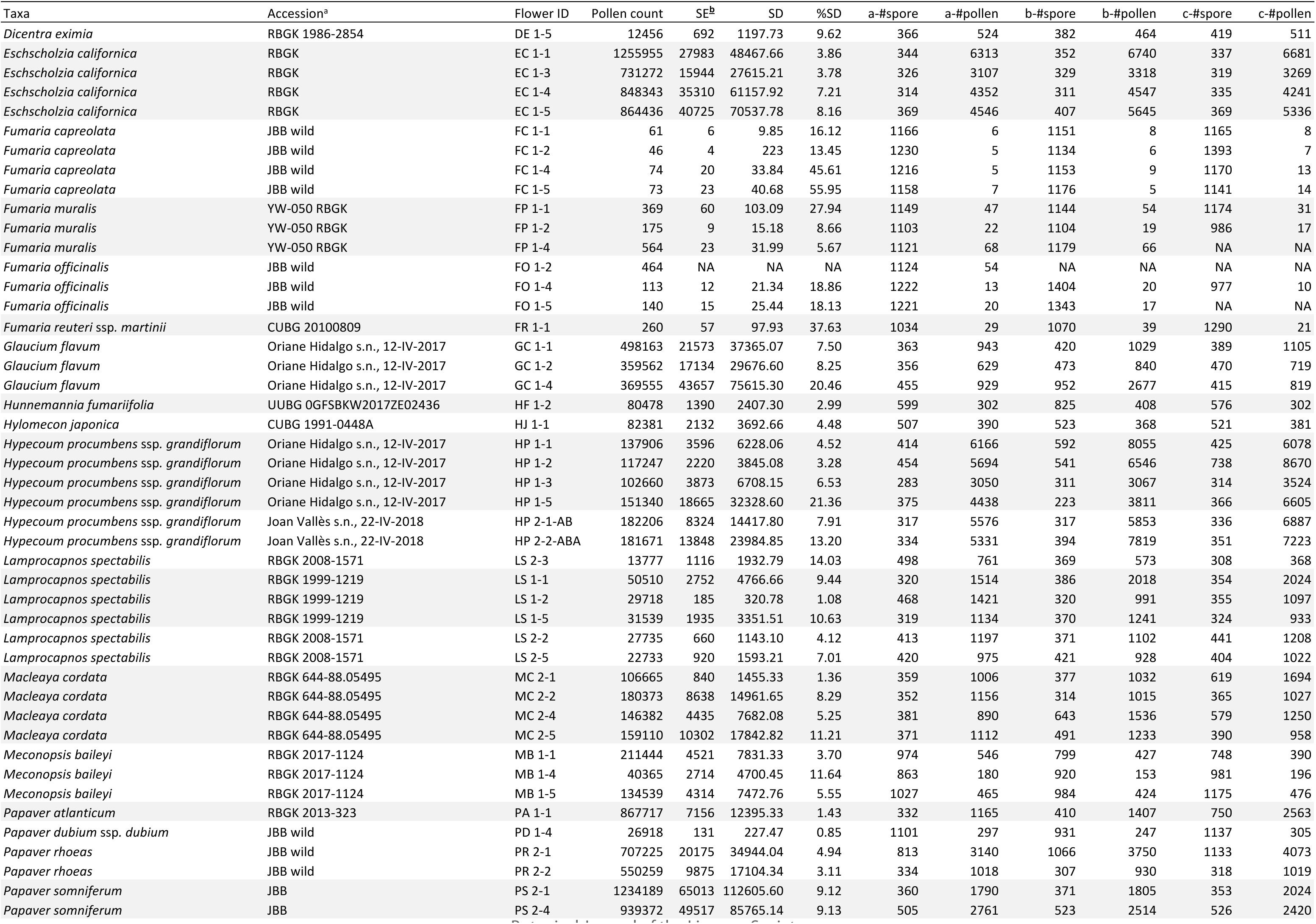
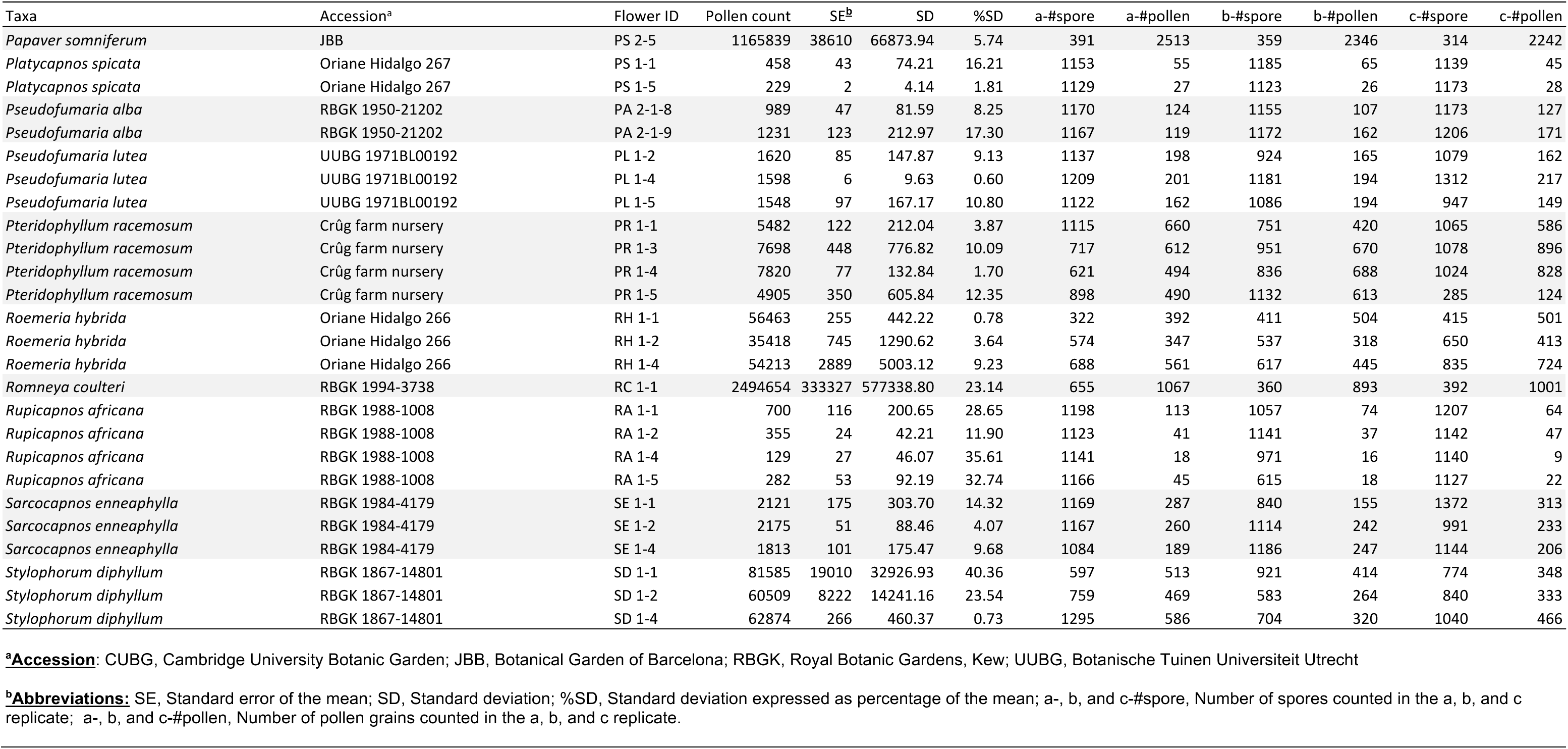
Total number of pollen grains per flower estimated for each species using flow cytometry. The sampling includes 38 species [only flow cytometry counts are available for *Corydalis temulifolia* Franch., *Cysticapnos pruinosa* (Bernh.) Lidén, *Platycapnos spicata* (L.) Bernh. and *Pseudofumaria alba* (Mill.) Lidén].

**Table S4.**
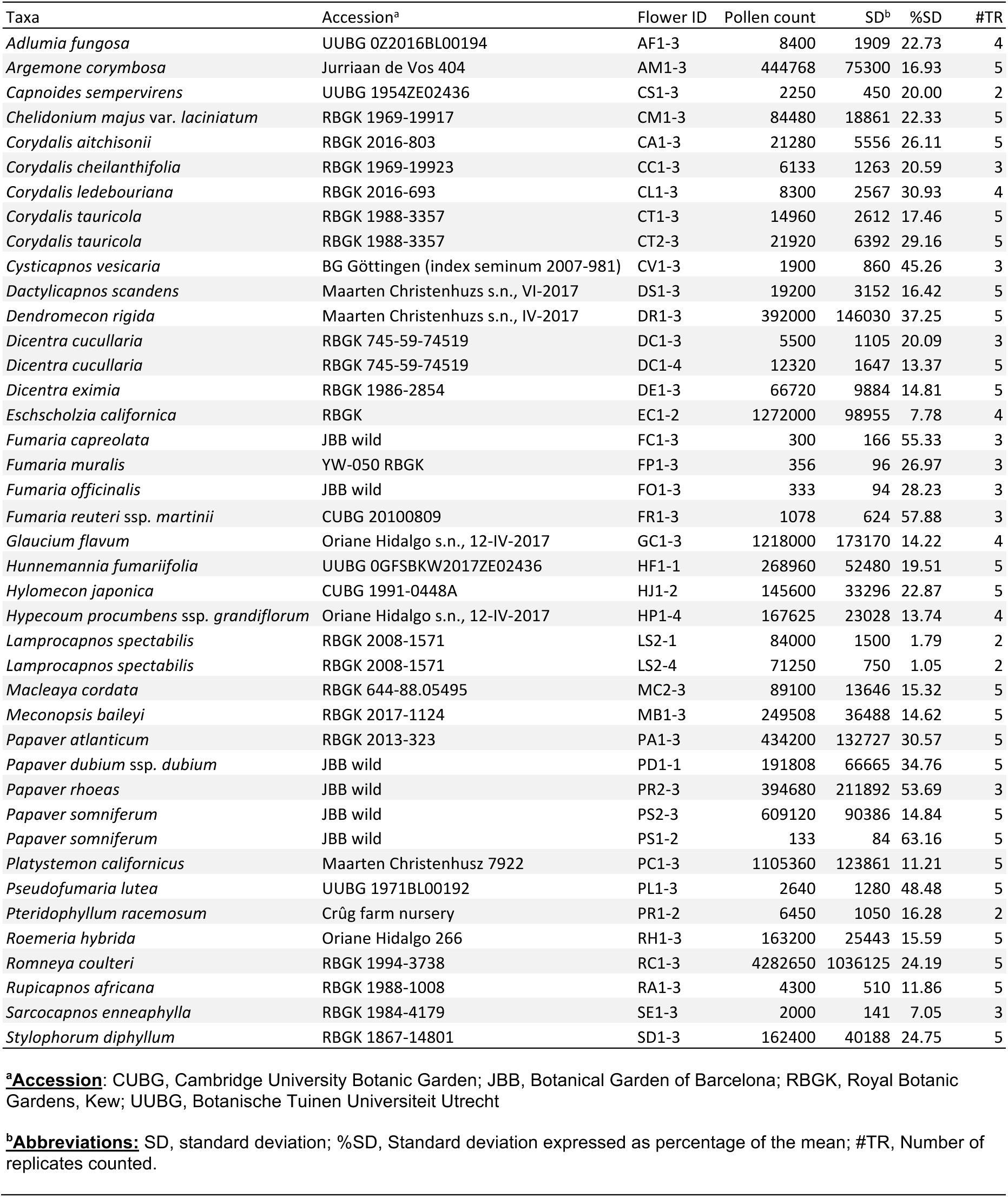
Total number of pollen grains per flower estimated for each species using manual counting. The sampling includes 37 species (only manual counts are available for *Corydalis* ledebouriana Kar. & Kir., *Dendromecon rigida* Benth. and *Platystemon californicus* Benth.).

